# Growth inhibitory effect of cerebrospinal fluid of multiple sclerosis patients on HFF2 cells

**DOI:** 10.1101/2025.09.20.671239

**Authors:** Sevda Jafari, Soheila Montazersaheb, Hamid Reza Heidari, Masoud Tabrizi-Nazhadyeh, Masoud Nikanfar, Mahnaz Talebi, Mohammad Yazdchi Marandi, Ata Mahmoodpoor, Narges Hejazi, Ommoleila Molavi, Mohammad Saeid Hejazi

## Abstract

Multiple sclerosis (MS) is an autoimmune disease characterized by demyelination of the central nervous system (CNS). Occurrence of immunogenic cell death (ICD) in this disease and expression of ICD markers in the CSF of MS patients are documented. However, cytotoxic effect of MS patients’ cerebrospinal fluid (CSF) on the cells is not reported. To evaluate the cytotoxic impact of CSF from MS patients on cell viability, human foreskin fibroblast (HFF2) cells were treated with CSF samples from individuals with MS and healthy controls. The results inferred that patient-derived CSF significantly declined the survival of HFF2 cells in comparison to the CSF of healthy individuals. This observation implies a cytotoxic environment and characteristic of patients’ CSF, which could be due to the presence of cellular-damaging factors such as calreticulin (CRT) in the patients’ CSF. The cytotoxic environment of CSF might reflect ICD occurrence in CNS of the patients and also could explain the progressive essence of the disease.

## Introduction

Multiple sclerosis (MS) is an autoimmune disorder affecting the central nervous system (CNS), marked by inflammation, demyelination, and progressive neurodegeneration (1, 2). Scientists have put forward two main ideas about how MS begins, which are known as the “outside-in” and “inside-out” theories (3, 4). The “outside-in” theory suggests that MS starts when inflammation in the body triggers the immune system to attack myelin. While the “inside-out” model indicates that death of primary CNS cells (e.g., oligodendrocytes) elicits subsequent autoimmunity against myelin (4, 5). Although the precise mechanisms underlying MS progression remain unclear, increasing evidence supports the involvement of regulated cell death pathways, particularly immunogenic cell death (ICD), in the disease’s pathogenesis (6, 7). ICD is a specific form of cell death that elicits a robust immune response through the release of danger-associated molecular patterns (DAMPs), such as high-mobility group box 1 (HMGB1), heat shock proteins (e.g., HSP70), calreticulin, ATP, and type I interferons like IFN-α (8–13). These molecules have been found at elevated levels in the cerebrospinal fluid (CSF) of MS patients, suggesting their role in aggravating neuroinflammatory responses (12). While our previous works documented the upregulation of several ICD markers in CSF of MS patients (14, 15), the functional implications of CSF composition on cellular viability remain underexplored. Therefore, we sought to investigate the direct effect of CSF from MS patients on cell viability using HFF2 cells as an in vitro model.

## Materials and Methods

### Cell Line and Sample Preparation

HFF2 cells were utilized as an *in vitro* model to evaluate the cytotoxic effects of patients’ CSF. The cells were cultured in DMEM with 10% FBS and 1% penicillin-streptomycin. To examine effect of CSF on cell viability, HFF2 cells were exposed to the pooled CSF samples obtained from three female MS patients (aged 30, 40, and 56 years) and two healthy individuals (one male, 37 years; one female, 61 years). Patients’ CSF samples were obtained from Nemooneh (Nomouneh) Clinical Laboratory in Tabriz, Iran. MS patients were diagnosed based on the revised McDonald criteria (16). Residual CSF samples were utilized, with no extra samples collected specifically for this project…

Control CSF samples were obtained from patients with isolated limb trauma, aged over 18 years old. The patients were admitted to the ICU of Shohada Orthopaedic Hospital and underwent surgery under spinal anesthesia. CSF sampling was carried out during spinal anesthesia after obtaining the patients’ informed consent. Patients with indwelling CNS devices or local/CNS infections were excluded. Both groups were matched based on ethnicity. All specimens were stored at −80 °C until being analyzed.

### MTT Assay for Cell Viability

MTT assay was conducted to assess the cytotoxic effects of patient-derived CSF on HFF2 cells. Due to limited sample volume, CSF from the MS patients and healthy controls were pooled into two separate groups prior to treatment.

HFF2 cells were seeded at a density of 7.5 × 10^3^ cells per well in a 96-well plate and cultured in 200 μl of culture medium containing 6% FBS and 30% CSF. Cells were kept for 48 hours at 37°C in a humidified atmosphere with 5% CO_2_. Following incubation, 20 μl of MTT solution (5 mg/ml) was added to each well, and the plate was further incubated for 4 hours to allow for formazan crystal formation. The medium was then carefully removed, and DMSO was added to dissolve the crystals. Absorbance was measured at 570 nm using a microplate reader (BioTek Instruments, Inc., USA). The experiment was carried out at least in triplicate. Data were analyzed using GraphPad Prism (version 8) and the results were expressed as the mean standard ± deviation (SD).

### Statistical analysis

The acquired data were analyzed using the GraphPad Prism software (version 8). Group comparisons were performed using one-way ANOVA, followed by Tukey’s post hoc test. Statistical significance was defined as a p-value of less than 0.05. All experiments were performed in triplicate.

## Results

To explore the potential cytotoxic effects of CSF from MS patients, we investigated its impact on cell viability using an in vitro model.

The results demonstrated a significant reduction in HFF2 cell viability following exposure to patient-derived CSF compared to that treated with control CSF. As depicted in Figure 1, the cell viability significantly decreased from 35.1 ± 2.11 % in the healthy group to 21.11 ± 3.21 % in the patient group (p<0.001). These results suggest that patients’ CSF exerts a cytotoxic/inhibitory effect on HFF2 cells. These findings align with the hypothesis that ICD-associated molecules present in MS patients’ CSF may contribute to neural damage by promoting stress responses and induction of regulated cell death in CNS-associated or exposed cell types. Interestingly, Pike and colleagues found that calreticulin (CRT) and its derivatives have a cytotoxic effect on cells.

**Figure 1.**
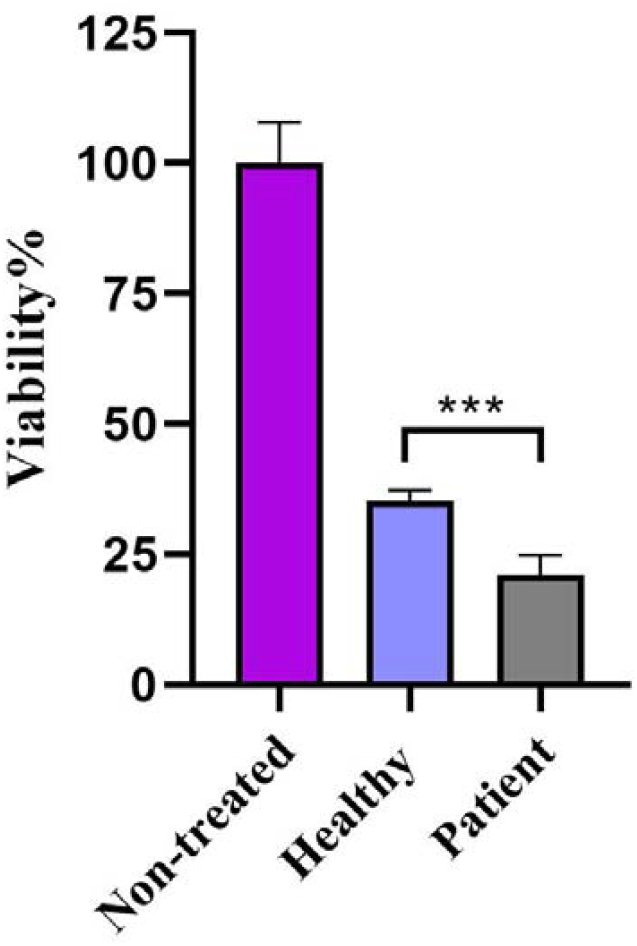
Cell Survival of HFF2 cells exposed to pooled CSF samples from MS patients and healthy controls. CSF from three MS patients and two healthy individuals was separately pooled before treatment of HFF2 cells. Cells were treated with the samples for 48 hours. The control group served as the nontreated cells. CSF of patients significantly reduced the cell viability as compared to the healthy group. Bar graphs are generated using GraphPad Prism software. The data represent the mean ± standard deviation (SD) from three independent experiments. ***p < 0.001.

Their study showed that CRT and its fragments are able to block the activity of endothelial cells and slow down tumor growth (17). Considering our paper reporting elevation of ICD markers including CRT in the CSF of MS patients (14, 15), the observed cell growth decline could also confer incidence of ICD and induction of related factors including CRT in CNS of the patients.

## Conclusion

This study demonstrates that CSF derived from MS patients significantly reduces the viability of HFF2 cells, indicating the cytotoxic potential of MS patients’ CSF. The observed cell growth inhibitory potential of the patients’ CSF suggests that factors present in the CSF may actively contribute to cellular damage beyond serving as biomarkers. These molecules, such as CRT, could result from the occurrence of ICD in CNS of the patients, and consequently could support the involvement of ICD in MS pathology. Additionally, our results might also explain the progressive nature of this disease and the beneficial effects of antioxidants as a complementary treatment in improving the condition of MS patients. Further studies with more samples are recommended.

## Funding

This research was funded by the Molecular Medicine Research Center at Tabriz University of Medical Sciences, Tabriz, Iran.

## Declaration of competing interest

The authors declare that they have no conflict of interest, financial or otherwise.

## Availability of data

The data sets used and/or analyzed during the current study are available from the corresponding author on reasonable request.

## Ethical Approval

The study received ethical consent from the Ethics Committee of Tabriz University of Medical Sciences, Tabriz, Iran (Ethics Code No: IR.TBZMED.REC.1403.218).

## Notes

### Competing Interest Statement

The authors have declared no competing interest.

